# Ribotoxic collisions on CAG expansions disrupt proteostasis and stress responses in Huntington’s Disease

**DOI:** 10.1101/2022.05.04.490528

**Authors:** Ranen Aviner, Ting-Ting Lee, Vincent B. Masto, Dan Gestaut, Kathy H. Li, Raul Andino, Judith Frydman

## Abstract

Huntington’s disease (HD) is a devastating neurodegenerative disorder caused by CAG trinucleotide repeat expansions encoding a polyglutamine (polyQ) tract in the Huntingtin (*HTT*) gene^1^. Although mutant HTT (mHTT) protein tends to aggregate, the exact causes of neurotoxicity in HD remain unclear^2^. Here we show that altered elongation kinetics on CAG expansions cause ribosome collisions that trigger ribotoxicity, proteotoxicity and maladaptive stress responses. CAG expansions cause an elongation rate conflict during HTT translation, when ribosomes rapidly decoding the optimal polyQ encounter a flanking slowly-decoded polyproline tract. The ensuing ribosome collisions lead to premature termination and release of aggregation-prone mHTT fragments. Due to the presence of a stress-responsive upstream open reading frame (uORF), HTT translation and aggregation are limited under normal conditions but enhanced under stress, seeding a vicious cycle of dysfunction. mHTT further exacerbates ribotoxicity by progressively sequestering eIF5A, a key regulator of translation elongation, polyamine metabolism and stress responses. eIF5A depletion in HD cells leads to widespread ribosome pausing on eIF5A-dependent sites, impaired cotranslational proteostasis, disrupted polyamine metabolism and maladaptive stress responses. Importantly, drugs that reduce translation initiation attenuate ribosome collisions and mitigate this escalating cascade of ribotoxic stress and dysfunction in HD.

## Main

HD is a progressive, age-dependent neurodegenerative disorder characterized by motor, cognitive and psychiatric symptoms^1^. It is caused by expansion of an uninterrupted CAG repeat segment in exon 1 of the HTT gene (herein HTT-ex1), encoding a variable-length polyQ tract. Repeats longer than 35 are pathogenic, and length is inversely correlated with age of onset^3^. CAG expansions in other genes cause similar neurodegenerative disorders (so-called “triplet disorders”), including spinocerebral ataxias^4^, suggesting common mechanisms of toxicity. It remains unclear how CAG expansion in a single gene can cause widespread dysfunction affecting multiple cellular pathways, including DNA repair, transcription, splicing, mitochondrial function, autophagy and stress responses^1–3^. The common hypothesis argues that toxicity is mediated by the mutant protein. Indeed, longer polyQ tracts tend to form oligomers, aggregates and inclusions associated with HD-like toxicity^1–3^. Both soluble oligomers and insoluble aggregates of mHTT can also sequester other proteins e.g. chaperones^5–7^, depleting them from the cellular pool and disrupting proteostasis, which is key to maintaining healthy cellular functions. However, formation of aggregates or inclusions can also be neuroprotective^8^, and neurons from affected individuals often lack aggregates^3^, suggesting a more complex relationship between CAG expansion and neurodegeneration.

An alternative hypothesis posits that the CAG repeat itself mediates RNA-based toxicity^9^. Thus, patients with pathogenic CAG expansions interspersed with the synonymous glutamine codon, CAA, exhibit delayed disease onset^10^. As these variants produce an identical polyQ tract, their protective effect cannot be due to changes in aggregation propensity of the mature protein. To reconcile these observations, we explored whether RNA-based toxicity in HD can result from aberrant translation dynamics of mHTT with an expanded CAG tract. CAG codons are rapidly decoded in mammals due to high abundance of their cognate and near-cognate (wobble) tRNAs^11^. The CAG tract may increase the risk of ribosome collisions due to an elongation rate conflict between fast and slow translating regions of the mRNA. Interestingly, previous studies showed the HTT mRNA harbors a uORF that can inhibit translation of a downstream reporter when ectopically expressed in cultured cells^12^. This uORF is evolutionarily conserved and has an optimal initiation context^13^ (Fig. 1a and Extended Data Fig. 1a). Since uORFs generally reduce translation initiation of downstream coding sequences (CDS)^14^, we hypothesized this uORF may reduce translation initiation on HTT mRNA and thus lower the risk of ribosome collisions, as reported for other transcripts^15,16^.

**Figure 1.**
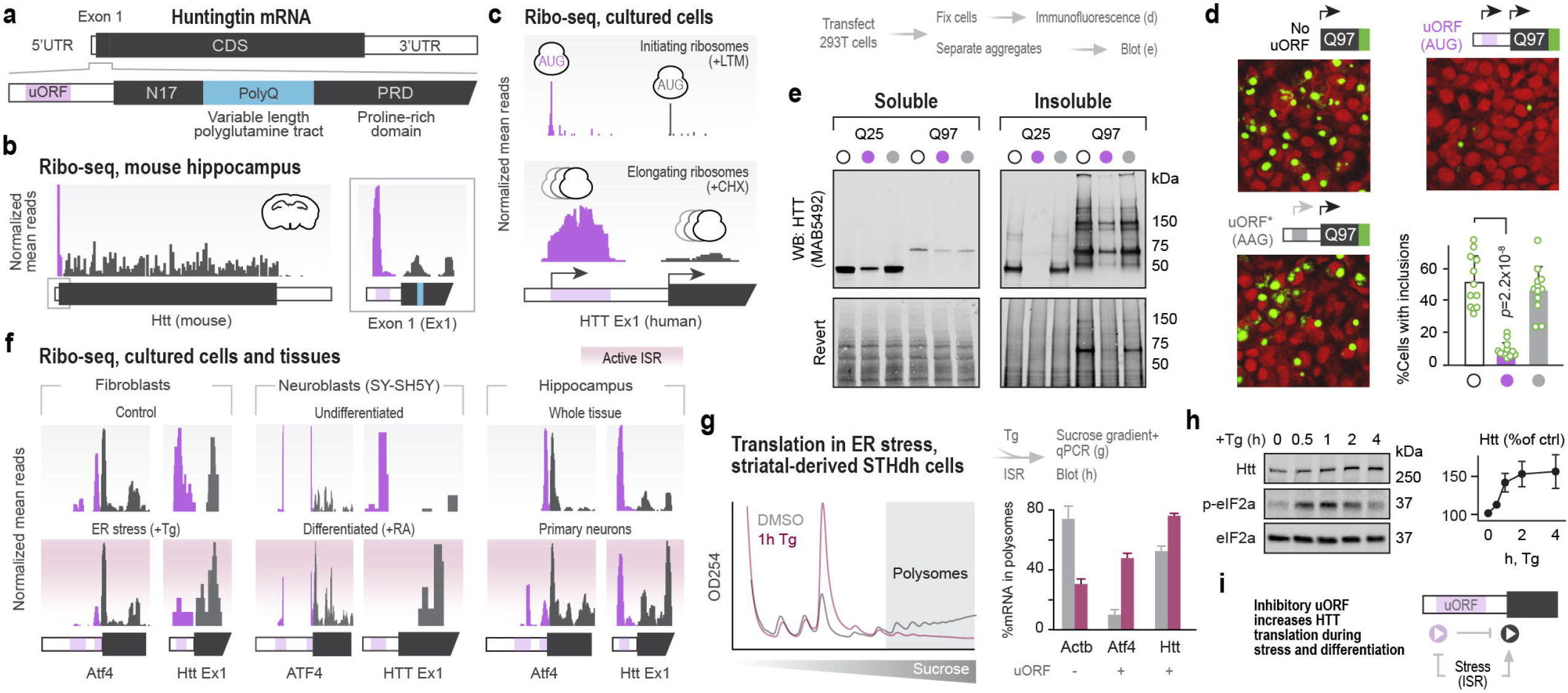
HTT expression is attenuated by a stress-responsive upstream open reading frame (uORF). **(a)** Schematic organization of human HTT mRNA. **(b-c)** HTT uORF is actively translated in vivo, inhibiting HTT expression. Ribosome profiling traces of elongating ribosomes on endogenous Htt mRNA in the mouse hippocampus (b, n=3), as well as traces of initiating (A-site positions, top) and elongating (footprints, bottom) ribosomes on the 5’ region of HTT in HCT116 cells (c, n=1). Pink, ribosome footprints on the uORF; gray, CDS. **(d-e)** HTT uORF reduces expression and aggregation of mHtt-ex1-GFP upon transfection in 293T cells. Aggregates were detected by either GFP fluorescence in fixed cells (d) or immunoblotting of insoluble fractions (e) at 24h post-transfection. Green, Htt; red, plasma membrane. Bar graph shows mean +/- s.d. of cells with puncta, counted manually from 15 random fields of each condition. *P*, p-value of a two-tailed Student t-test. Blots are representative of 3 independent repeats. **(f)** uORF skipping occurs on both Htt and Atf4 increases under ISR conditions, including thapsigargin (Tg)-induced ER stress in mouse fibroblasts (n=2), retinoic acid (RA) differentiation of SH-SY5Y neuroblasts (n=2) and cultured hippocampal tissues (n=1). **(g-h)** Translation of endogenous Htt increases in response to ER stress in wild-type (Q7/7) mouse striatal cells. Cells were treated with 1 μM Tg for 1h (g) or up to 4h (h) and subjected to qPCR of sucrose gradients or immunoblots, respectively. Shown are means +/- s.d. of 3 independent repeats for qPCRs and representative blots and densitometry from 3 independent repeats. (i). Proposed model.

Using a combination of cellular and animal HD models, we tested these hypotheses and found that the HTT uORF plays a protective role by attenuating translation initiation, whereas CAG expansions increase the risk of ribosome collisions, resulting in higher rates of premature termination of nascent mHTT. Strikingly, mHTT protein also plays a role in translation dysfunction, as it sequesters translation factor eIF5A and thus promotes widespread ribosome pausing on hundreds of transcripts, leading to widespread proteostasis collapse. Importantly, we show the ensuing ribotoxicity and proteotoxicity in HD can be counteracted using a small molecule inhibitor of translation initiation.

### A stress-responsive uORF attenuates HTT translation and aggregation

To determine how the uORF affects endogenous HTT translation, we analyzed a panel of ribosome profiling datasets obtained from human and mouse cells and tissues. Consistent with the uORF inhibiting HTT translation in the central nervous system, we observed higher ribosome density on the uORF than the downstream CDS of HTT in mouse hippocampus^17^ (Fig. 1b). Treatment of cultured human cells with either lactimidomycin (LTM) or harringtonine (Harr.)^18^, which arrest ribosomes at AUG codons used as translation start sites, confirmed that initiation occurs separately on both the uORF and CDS, with elongating ribosomes occupying the entire length of the uORF but not flanking regions (Fig. 1c, Extended Data Fig. 1b). These observations suggest that the endogenous uORF inhibits translation of HTT *in vivo* by lowering initiation on the CDS. Because HTT aggregation is concentration dependent^3^, we tested whether the presence of a functional uORF in the HTT mRNA reduces soluble and insoluble HTT protein levels. Expression of fluorescently-tagged HTT-ex1 variants with and without the endogenous 5’ untranslated region (UTR) of human *HTT* confirmed that the uORF reduces protein expression and aggregation in transfected 293T cells (Fig. 1d-e and Extended Data Fig. 1c). This protective effect was lost by mutating the upstream AUG to AAG, which renders the uORF non-functional (Fig. 1d-e). These experiments indicate that active initiation on the uORF attenuates HTT translation and aggregation.

Translation of uORF-regulated polypeptides is known to respond to conditions that induce eIF2a phosphorylation e.g. stress or differentiation^19^. Induction of the integrated stress response (ISR) leads to increased translation of the main ORF through a mechanism that involves eIF2a phosphorylation and uORF bypassing^19^. One well-characterized example is ATF4, the main effector of the ISR and a transcription factor actively translated during stress to promote recovery^19^. To determine whether stress increases uORF bypassing on the endogenous HTT mRNA, we analyzed ribosome profiling datasets collected under conditions of ER stress (thapsigargin, Tg) or neuronal differentiation (retinoic acid, RA). Fibroblasts^20^ and neuroblasts^21^ responded to stress and differentiation by increasing uORF bypassing on both ATF4 and HTT (Fig. 1f). Interestingly, analysis of a broad panel of tissues, cell types and conditions revealed dramatic differences in the relative ribosome densities on HTT uORF vs main ORF, indicating that HTT translation can vary widely (Extended Data Fig 1d). For example, we observed lower ribosome density on HTT uORF in primary hippocampal neurons as compared to the parental tissue from which they were obtained^8^ (Fig. 1f). These observations may be relevant to assess the impact of stress and differentiation on HTT protein levels, which are linked to HD vulnerability.

To extend and validate these observations, we examined the effect of ER stress on HTT translation and protein levels in neuron-like cells isolated from wild-type mouse striatum^22^ (hereinafter referred to as “striatal cells”). We induced ER stress with Tg and measured the association of Htt mRNA with translating polysomes by sucrose gradient fractionation coupled to qPCR. Tg treatment reduced global translation rates and displaced the housekeeping beta-actin (Actb) mRNA from polysomes (Fig. 1g and Extended Data Fig. 1e), consistent with attenuated translation during stress. In contrast, Atf4 and Htt mRNAs responded to Tg treatment by shifting deeper into polysomes (Fig. 1g and Extended Data Fig. 1e), as expected for uORF-regulated transcripts whose translation is induced during the ISR. Importantly, immunoblot analysis confirmed that Htt protein levels increase during ER stress (Fig. 1h). Taken together, these findings indicate that HTT translation is dynamically regulated by its uORF across a wide range of cell types and stress conditions (Fig. 1i).

### CAG expansions promote ribosome collisions on mHtt mRNA

Given that the uORF reduces initiation—and therefore ribosome density—on HTT CDS, we explored how varying initiation rates and CAG repeat lengths might affect HTT translation dynamics. We employed RFMapp^23^, a ribosome flow model that uses the tRNA adaptation index (tAI) as a proxy for elongation rates, to simulate ribosome distribution on HTT-ex1. Due to differences in codon optimality^11^, ribosomes are predicted to elongate quickly through CAG repeats and slow down on rare CCG proline codons in the downstream proline rich domain (PRD, Fig. 1a). Of note, translation of multiple consecutive prolines is further slowed by their physicochemical properties^24^, suggesting our simulation may underestimate the ribosome dwell time on the PRD. At an average rate of initiation (0.05/sec)^25,26^, the model predicts a CAG length-dependent buildup of ribosomes upstream of the PRD, with stacking of three or more ribosomes on pathogenic length CAG expansions (Fig. 2a). A 100-fold reduction in initiation rate, but not 50-fold reduction, completely prevented ribosome buildup (Fig. 2a). This simulation suggests that CAG expansions increase the risk of ribosome collisions and that such risk can be mitigated by reduced initiation rates.

**Figure 2.**
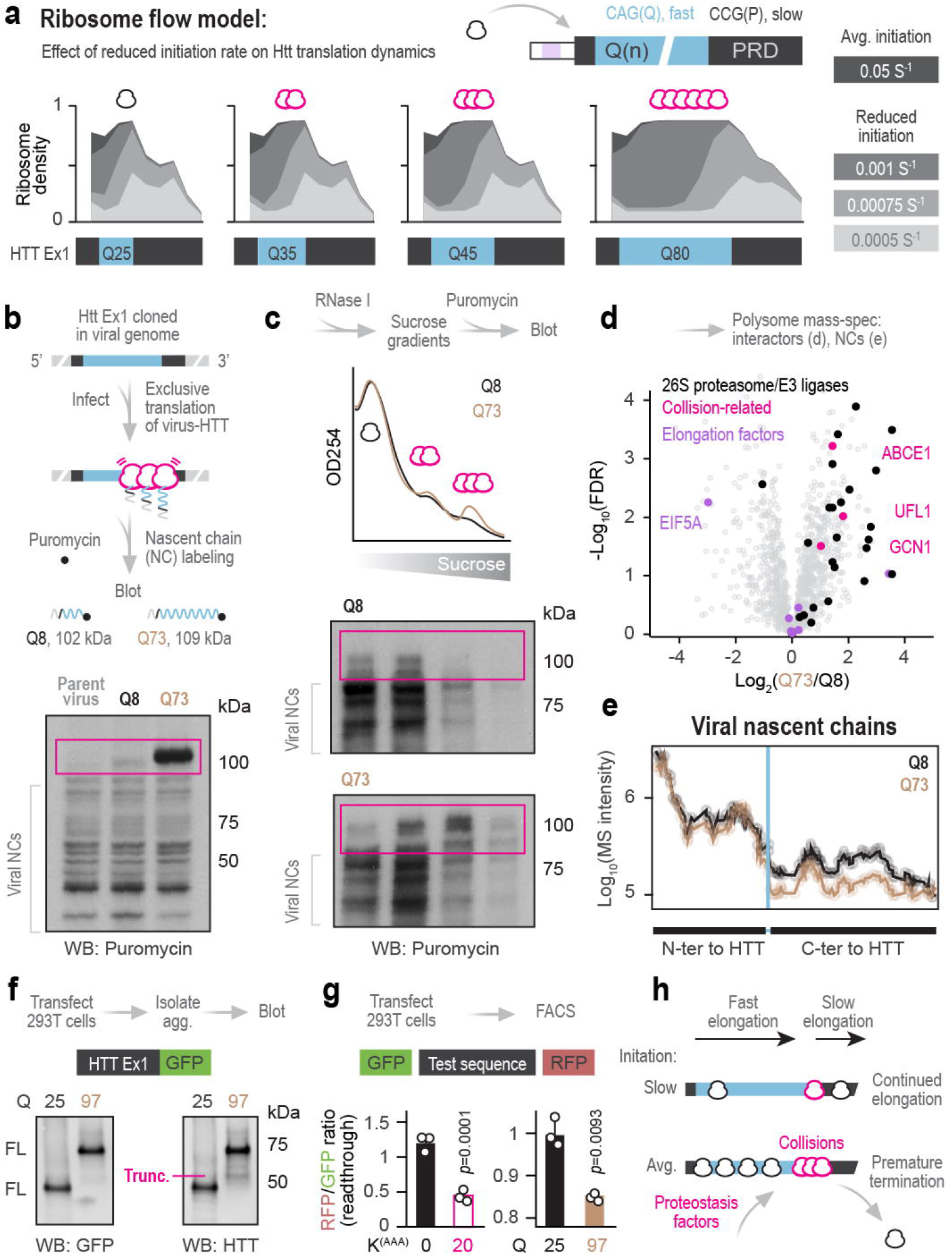
Ribosomes collide on CAG expansions and terminate prematurely. **(a)** Translation of mHtt is associated with higher risk of ribosome collisions due to rapid decoding of CAGs in the polyQ tract and slower decoding of CCGs in the proline-rich domain (PRD). Shown are ribosome flow model predictions of ribosome density on wild-type (Q25, Q35) and mutant (Q45, Q80) HTT, using *Homo sapiens* tAI, chunk size=10 and the indicated initiation rates. **(b)** Ribosomes accumulate on mHtt-ex1. Wild-type (Q8) or mutant (Q73) Htt-ex1 was cloned into poliovirus genome. Huh7 cells were infected with the engineered viruses, inducing rapid shutoff of mRNA translation and homogenous, synchronized translation of Htt as part of the viral polypeptide. At 3.5 hours, cells were treated with puromycin to label nascent polypeptide chains (NCs) and analyzed by immunoblotting (n=5). Purple box indicates virus-Htt polyproteins prior to cleavage by the viral protease. **(c)** Translation of mHtt involves ribosome collisions. Huh7 cells were infected as before, lysed and treated with RNase I. Lysates were fractionated on 10-50% sucrose gradients and fractions were incubated with puromycin. Top, polysome profiles; bottom, immunoblot (n=3). **(d-e)** Ribosome collisions on mHtt are associated with recruitment of quality control factors and premature termination. Huh7 cells were infected as before, lysed and fractionated on 10-50% sucrose gradients. Polysome fractions were pooled, pelleted through a sucrose cushion and analyzed by mass-spectrometry (MS; n=3). Shown are pairwise comparisons of individual polysome-associated proteins (d) and MS intensities of viral peptides mapping N- or C-terminally to the Htt insert (e). **(f-g)** Translation of mHTT-ex1 is associated with increased premature termination in striatal and 293T cells. Immunoblot analysis of insoluble fractions from 293T cells transfected with HTT-ex1-GFP for 24h. Shown are representative blots from 3 independent repeats (f). Flow cytometry analysis of 293T cells transfected with plasmids encoding GFP and RFP from a single ORF, separated by a linker. The ratio of RFP to GFP is inversely proportional to the rate of premature termination. K^(AAA)^0/20, linker with 0 or 20 consecutive repeats of AAA, known to strongly induce premature termination. Shown are means +/- s.d. of 3 independent repeats. *P*, p-value of a Student t-test (g). **(h)** Proposed model.

Testing these predictions on endogenous HTT mRNA was complicated by its low abundance and the inability to accurately map repetitive elements by ribosome profiling. Cell free translation, on the other hand, does not reflect the initiation and elongation kinetics and tRNA component of intact cells. We thus developed a virus-based system to express, label and detect HTT nascent chains (NCs) in cells. We introduced the coding sequence for wild-type polyQ(8) or mutant polyQ(73) human HTT-ex1 into the single ORF of poliovirus, flanked by sites for viral protease cleavage. Upon infection, poliovirus rapidly shuts off cellular mRNA translation and reprograms ribosomes to exclusively translate viral RNA from an internal ribosome entry site^27^. In this system, HTT-ex1 is translated as part of the viral polyprotein, leading to the generation of ribosome-bound HTT-ex1 nascent chains. Consistent with a general biosynthetic defect caused by polyQ expansion, we find that viruses harboring mutant polyQ(73) replicate more slowly than their wild-type polyQ(8) counterparts (Extended Data Fig. 1a). To monitor ribosome buildup on transcripts encoding HTT-ex1, we labeled NCs with puromycin, which becomes covalently incorporated by elongating ribosomes and can be detected by puromycin-specific immunobloting^28^. If elongation rates are uniform and ribosomes are evenly distributed on the RNA template, the size of puromycylated NCs will also be evenly distributed; in contrast, ribosome buildup at a specific region of the RNA will generate puromycylated NCs of a discrete size. As predicted by the RFMapp model, introduction of mutant but not wild-type HTT-ex1 produced a strong, clearly defined puromycylated band indicative of stalled ribosomes, when expressed in either Huh7 (Fig. 2b and Extended Data Fig. 2b) or SH-SY5Y (Extended Data Fig. 2c) cells. The size of this puromycylated NC product is consistent with ribosome buildup at the end of the expanded CAG repeat region, prior to cleavage of N-terminal viral sequences by the viral protease.

Collided ribosomes form a tight interface that is resistant to RNase digestion^29^. If the defined polyQ(73)-dependent puromycylated NC is produced by ribosome collisions on mHTT, it should be associated with RNase-resistant disomes or higher order species. To test this prediction, we infected Huh7 cells as above and digested lysates with RNase I followed by sucrose gradient fractionation to separate monosomes from RNAse-resistant polysomes. The presence and size of NCs in each fraction was assessed by puromycin labeling. Consistent with an induction of ribosome collisions, translation of polyQ(73) led to an increase in overall levels of RNAse-resistant trisomes as compared to polyQ(8) (Fig. 2c, top). Importantly, ribosomes stalled on polyQ(73) NCs, but not polyQ(8) or parent virus alone, were predominantly associated with RNAse-resistant trisomes (Fig. 2c and Extended Data Fig. 2d).

Collided ribosomes are known to recruit specific proteostasis factors to promote either resolution or premature termination and clearance^30–32^. To better understand how the presence of polyQ(73) remodels translating polysomes, we next examined the overall composition of polyQ(8) vs polyQ(73) polysomes by mass-spectrometry. We isolated polysomes from infected cells (Extended Data Fig. 2e) and analyzed their interacting proteins by mass-spectrometry (MS). Translation of polyQ(73) was associated with changes in ribosome interactors, most notably increased recruitment of ubiquitin-proteasome system (UPS) components, including proteasomal subunits and E3 ubiquitin ligases, as well as collision-sensing ribosome quality control (RQC) factors e.g. GCN1 (Fig. 2d and Supplementary table 1a).

We next tested whether ribosome collisions on polyQ(73) increase premature translation termination, which would lead to release of mHtt-ex1 fragments. If this were the case, fewer ribosomes should be translating the viral sequences downstream of the mHTT insert. Indeed, proteomic analysis of polysomes revealed a reduction in the levels of viral nascent chains downstream of polyQ(73) as compared to polyQ(8) (Fig. 2e). Ribosome profiling analyses revealed a similar reduction in ribosome occupancy downstream of polyQ(73) (Extended Data Fig. 2f).

To confirm that CAG expansions in mHtt-ex1 promote abortive ribosome collisions, we transfected 293T cells with mHTT-ex1-GFP and analyzed their insoluble fraction. If ribosomes terminate prematurely, the GFP moiety should be lost; indeed, immunoblot analysis detected the presence of truncated polyQ(97) but not polyQ(25) lacking the downstream GFP (Fig. 2f). In addition, we adapted a previously established reporter of collision-induced premature translation termination, consisting of a transcript encoding GFP-K^AAA^20-mCherry. Previous studies showed that when ribosomes collide and terminate on the K^AAA^20 stalling motif, which mimics PolyA tail translation in defective nonstop mRNAs, the ratio of mCherry to GFP produced is decreased^33^. We thus replaced the K^AAA^20 stalling motif in this reporter for HTT-ex1 polyQ(25), which should not stall translation, or for polyQ(97), which should. Both K^AAA^20 and mutant HTT-ex1 polyQ(97) reduced the ratio of mCherry to GFP as compared to K^AAA^0 or polyQ(25) control (Fig. 2g). Taken together, our combined modelling and experimental approaches shows that CAG expansions in HTT-ex1 increase the risk of ribosome collisions, triggering recruitment of proteostasis and RQC factors, and resulting in premature termination and release of N-terminal mHTT-ex1 fragments, known to be neurotoxic^1^ (Fig. 2h).

### PolyQ-expanded mHtt depletes eIF5A from polysomes and disrupts its function in translation

In addition to the selective recruitment of RQC and UPS components, our proteomic analysis revealed a depletion of translation factor eIF5A from polysomes translating mHTT-ex1 (Fig. 2d). eIF5A plays multiple critical roles in translation initiation, elongation and termination, and regulates polyamine metabolism and stress responses^24,34^. To determine how mHTT could prevent eIF5A from associating with polysomes, we analyzed a panel of proteomic datasets generated using various cellular and mouse models of HD. Analyses of HTT-ex1 expressed in either yeast^35^ or mouse neuroblasts^36^ indicate that soluble mHTT binds eIF5A with higher affinity than wild-type controls (Fig. 3a). Additionally, eIF5A was enriched in aggregates from neuroblastoma cells expressing mHTT-ex1^37^ (Fig. 3b). We also examined a longitudinal dataset comparing the soluble and aggregated brain proteomes of pre-symptomatic and post-symptomatic R6/2 HD mice and wild-type age-matched controls^6^. R6/2 mice express mHtt-ex1 with polyQ(>140) leading to symptom onset around 8 weeks of age. Consistent with the findings in cultured cells, eIF5A was highly enriched in the insoluble fraction of aged but not young asymptomatic animals (Fig. 3c). Analysis of soluble eIF5A levels revealed an inverse correlation with the levels of soluble mHtt, which was restricted to aged R6/2 mice (Fig. 3d). Interestingly, the mHtt-dependent reduction of soluble eIF5A in aged HD mice was preceded by an earlier increase in soluble eIF5A at 8 weeks (Extended Data Fig. 3a-b). Levels of other factors involved in sensing and resolving ribosome collisions were also higher at 8 weeks (Extended Data Fig. 3a-b), suggesting an early adaptive response to a higher burden of collisions in HD mice. Importantly, analysis of brain proteomes from another HD mouse model, zQ175^38^, showed a similar relationship between mHtt and eIF5A in both soluble (Fig. 3e) and insoluble (Extended Data Fig. 3c) fractions.

**Figure 3.**
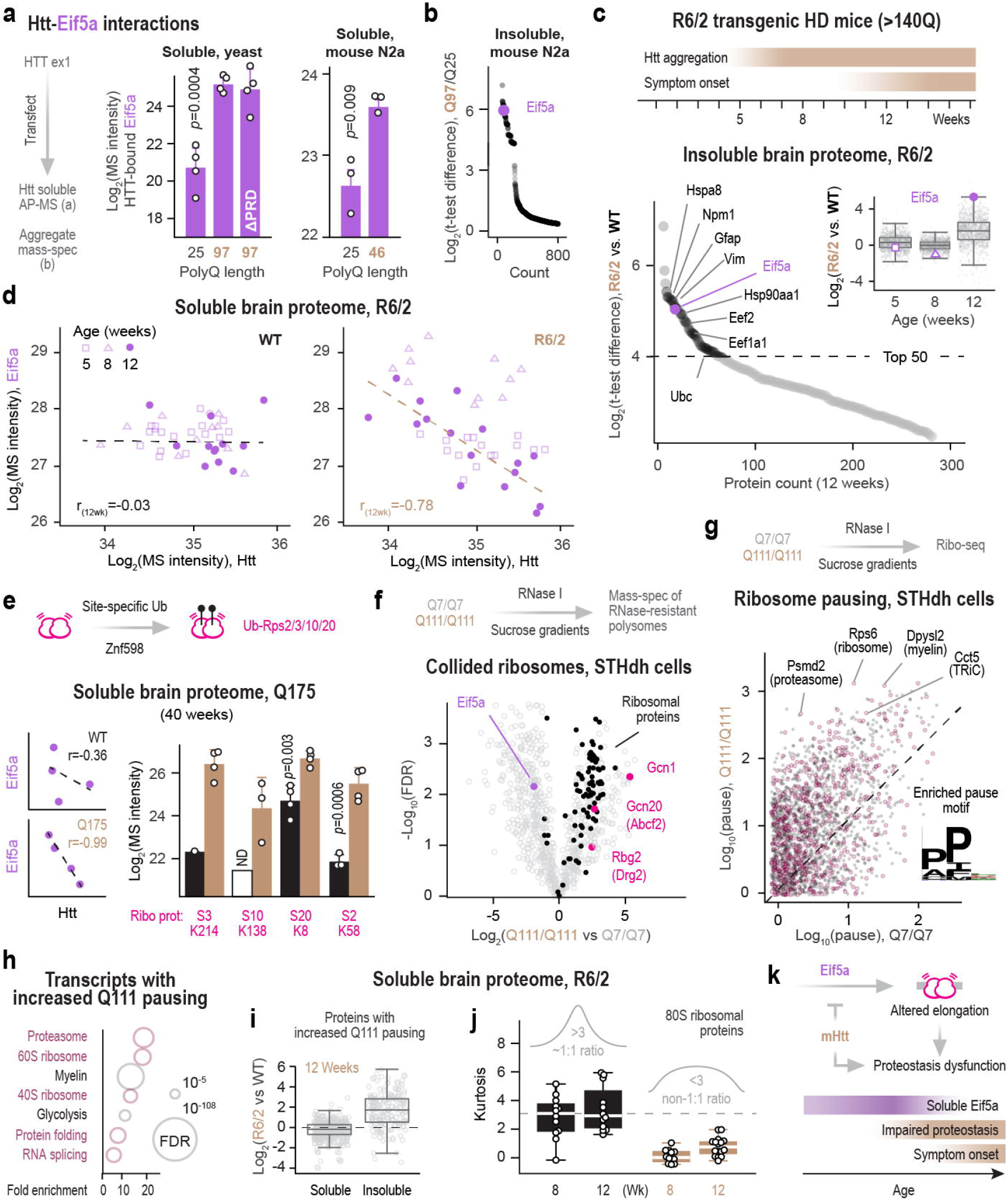
mHtt disrupts global translation dynamics by sequestering translation factor eIF5A. **(a)** PolyQ expansions increase affinity of eIF5A to mHTT. Yeast (left) and mouse neuroblastoma cells (right) were transfected with wild-type or mutant HTT-ex1. Following affinity purification, soluble HTT-ex1 interactors were detected by MS (n=4 and 3 for yeast and N2a, respectively). **(b)** Eif5a is highly enriched in the insoluble fraction of N2a cells transfected with mHTT-ex1 (n=4). **(c)** Eif5a is highly enriched in the insoluble fraction of brains from aged but not young R6/2 mice. Top, timeline of mHtt aggregation and symptom onset in the transgenic R6/2 mouse HD model. Bottom, t-test difference in proteins detected by MS using the insoluble brain proteome of aged (12 weeks) R6/2 versus wild-type mice. Inset, same analysis for early, intermediate and late ages (5, 8 and 12 weeks, respectively). n=12 and 16 for wildtype and R6/2, respectively. Boxplots show medians with upper and lower quartiles, and whiskers are 1.58xIQR. **(d)** Eif5a levels are inversely correlated with levels of mutant but not wild-type Htt, and only in aged mice. Pairwise comparisons of eIF5A and Htt levels in the soluble brain proteome of wild-type and R6/2 mice (n=12/16). **(e)** Ubiquitination of ribosomal proteins in the brains of aged HD mice is consistent with increased ribosome collisions. Left, pairwise comparisons of eIF5A and Htt in 40-week-old wild-type and Q175 mice. Right, relative ubiquitination of specific sites on ribosomal proteins, as determined by MS analysis (n=4). **(f)** Collided ribosomes in HD striatal cells are depleted of eIF5A. Mouse striatal cells expressing either polyQ(7) or polyQ(111) Htt were lysed, digested with RNase I and fractionated on sucrose gradients. MS analysis was performed on fractions of the gradient harboring collided, RNase-resistant ribosomes (n=3). **(g)** Ribosome pausing increases in HD striatal cells, particularly on eIF5A-dependent sequence motifs. Shown is a pairwise comparison of pause scores calculated using ribo-seq data from striatal cells as in (f). **(h)** Fisher enrichment analysis of Gene Ontology functions enriched among pause-containing transcripts from (g). **(i)** Proteins encoded by pause-containing transcripts are enriched in the insoluble brain proteome of aged 12-week-old R6/2 mice (n=12/16). **(j)** Ribosomal protein stoichiometry is disrupted as early as 8 weeks of age in R6/2 brains. Shown are kurtosis scores for all core ribosomal proteins as measured by MS (n=12/16). **(k)** Proposed model.

We next examined the consequences of eIF5A depletion in HD models. Proteomic analysis of ubiquitinated soluble brain proteins from aged zQ175 mice revealed increased ubiquitination of 40S ribosomal proteins on sites associated with ribosome collisions^39^ (Fig. 3e) but not other sites (Extended Data Fig. 3d), suggesting a persistent defect in translation elongation dynamics. To better define how loss of eIF5A affects translation and cellular responses in HD, we employed a well-established cell model of striatal neuron-like cells from mice expressing two copies of full-length Htt with either polyQ(7) or polyQ(111)^22^. First, we used sucrose gradient purification followed by MS analyses to compare the composition of RNase-resistant polysomes from polyQ(7) or polyQ(111) cells. Similar to our viral mHtt-ex1(Q73) (Fig. 2d), expression of an expanded polyQ tract increased polysome association of the Gcn1-Gcn20-Rbg2 complex, which binds collided ribosomes^32^. Importantly, we also observed decreased association of eIF5A in the polyQ(111) cells (Fig. 3f and Supplementary table 1b). We next subjected these cells to ribosome profiling analyses and detected an overall increase in ribosome pausing in the polyQ(111) cell line (Fig. 3g), as previously reported^40^. Sequence motif analysis confirmed that HD-linked pause sites are enriched for PPX, a known eIF5A-dependent motif (Fig. 3g). Of note, increased pausing disproportionately affects mRNAs encoding for proteins involved in proteostasis, including ribosomal and proteasomal proteins, as well as the chaperonin TRiC/CCT and factors involved in myelination and glycolysis (Fig. 3g-h, Extended Data Fig. 3e and Supplementary table 2). Consistent with observations that aberrant pausing can lead to aggregation of stalled nascent polypeptides^41^, the protein products of mRNAs showing increased collisions were enriched in the insoluble brain fraction of aged R6/2 mice (Fig. 3i). Analysis of the soluble brain proteome of R6/2 mice was consistent with defects in the biogenesis of these proteostasis machines. For instance, we found evidence for an early loss of 1:1 stoichiometry among the core ribosomal and proteasomal proteins in the brains of R6/2 mice (Fig. 3j and Extended Data Fig. 3f). Thus, the increase in aberrant translation pausing on mRNAs encoding key proteostasis complexes may impact their biogenesis, seeding a vicious cycle of proteostasis dysfunction that contributes to HD symptoms. Indeed, mutations affecting ribosomal and proteasomal stoichiometry are linked to neurodegeneration in mice^42^.

Taken together, we find that mHTT-ex1 protein sequesters eIF5A away from polysomes, altering translation elongation dynamics across hundreds of mRNAs and resulting in proteostasis defects. In HD mice, symptom onset is associated with eIF5A depletion from the soluble proteome, likely fueled by aggregation. This is preceded by an early biosynthetic disruption of major proteostasis machineries and an early compensatory increase in soluble eIF5A and RQC factors (Fig. 3k).

### eIF5A-linked impairment of polyamine metabolism and stress responses in HD

By facilitating translation elongation on ribosome pause sites e.g. PPX, eIF5A supports protein and organelle biogenesis, autophagy and immune functions^43^—all of which are key cellular pathways often deregulated in HD^1,3^. eIF5A is also involved in the regulation of key enzymes mediating polyamine metabolism^44^. Furthermore, yeast experiments also implicate eIF5A in the uORF bypassing required for translational induction of ISR effectors e.g. ATF4 and GADD34/PPP1R15A^45,46^ (Fig. 4a). We thus explored whether mHtt expression in HD cells leads to dysregulation of these processes. Using ribosome profiling data from wild-type and HD striatal cells, we calculated the ratio of ribosomes on uORFs vs CDS for individual mRNAs across the transcriptome. Striatal cells expressing polyQ(111) exhibited significant differences in uORF-to-CDS ratios compared to polyQ(7), consistent with changes in translation regulation of a subset of transcripts (Fig. 4b). Of note, these included both ISR effectors and polyamine-regulating enzymes (Fig. 4b). Under normal conditions, translation of Azin1, Amd1/2 and Sat1 is co-regulated by polyamine levels^47^, but in HD polyQ(111) cells their co-regulation is lost (Fig. 4b-c). Direct measurement of polyamine levels in these HD cells confirmed a reduction in polyamine levels (Fig. 4d).

**Figure 4.**
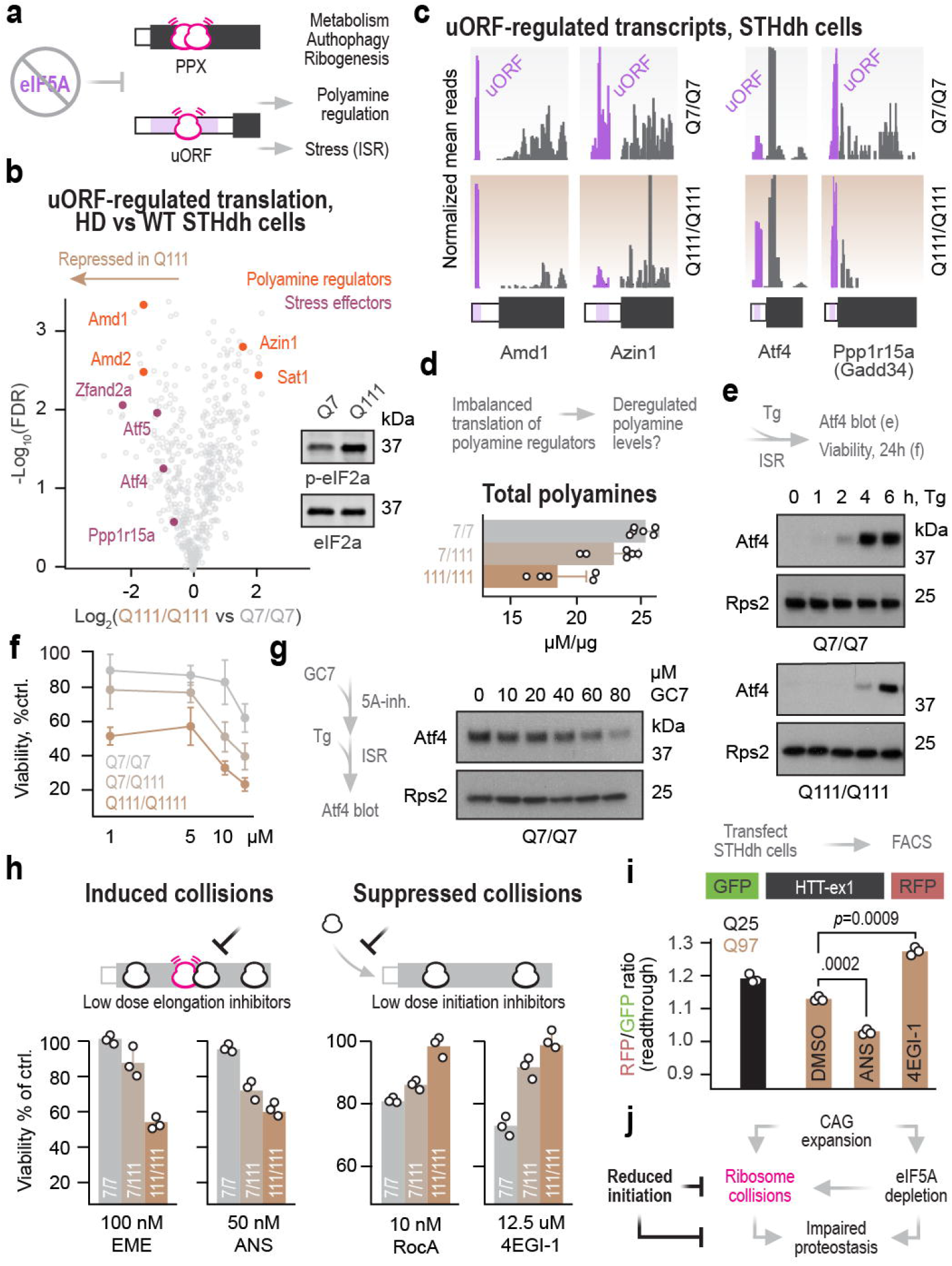
Modulating translation initiation mitigates harmful effects of eIF5A loss on elongation kinetics. **(a)** eIF5A not only promotes translation elongation on transcripts important for multiple cellular pathways; it also regulates translation initiation on uORF-containing transcripts. **(b)** HD striatal cells show changes in translation of uORF-regulated transcripts. Pairwise comparison of uORF repression, calculated at the ratio of ribosomes on the uORF vs CDS of individual transcripts across the transcriptome. The left of the plot reflects attenuated translation in polyQ(111)-expressing cells (n=3). Inset, immunoblot of resting striatal cells (n=2). **(c)** Ribosome profiling traces of transcripts highlighted in (b). **(d)** Quantification of total polyamine levels in HD striatal cells (n=6). **(e)** Induction of neuroprotective ISR is delayed in HD striatal cells treated with Tg (n=3). **(f)** HD striatal cells are hypersensitive to Tg, as determined by ATP measurements at 24h of treatment with the indicated concentrations (n=3). **(g)** eIF5A is required for ISR induction. Wild-type striatal cells were treated for 48h with increasing concentrations of GC7, an eIF5A inhibitor. Tg was added for additional 4h to induce the ISR (n=3). **(h)** HD striatal cells are hypersensitized to inhibitors of elongation and desensitized to inhibitors of initiation, as determined by ATP measurements at 24h of treatment with the indicated drugs (n=3). **(i)** 4EGI-1 improves translational readthrough on mHTT-ex1. Wild-type striatal neurons were transfected with plasmids encoding for GFP and RFP, as in Figure 2f. The indicated drugs were added at 6 hours and flow cytometry was performed at 24 hours post-transfection (n=3). (j) Proposed model.

We also examined the regulation of ISR genes in HD cells. mHtt expression can trigger eIF2a phosphorylation through multiple mechanisms, including the unfolded protein response^48^. Indeed, cells expressing polyQ(111) show increased basal phosphorylation of eIF2a (Fig. 4b) and reduced global translation rates (Extended Data Figure 4a), both hallmarks of ISR activation. Although eIF2a phosphorylation should increase uORF bypassing^19^, cells expressing polyQ(111) had higher ribosome densities on the uORFs of major ISR effectors Atf4, Atf5 and Gadd34 (Ppp1r15a) (Fig. 4b-c and Extended Data Figure 4b). A similar trend was observed for Htt itself (Extended Data Figure 4b-d), consistent with its uORF-dependent regulation and the co-regulation of Atf4 and Htt (Figure 1). To directly test whether the reduced uORF bypass on ISR effectors in HD cells leads to dampening of the translational response to stress, we induced acute ER stress in wild-type and HD striatal cells and measured both Atf4 levels and cell viability. Cells expressing polyQ(111) showed a delay in Atf4 expression (Fig. 4e) and were hypersensitive to Tg (Fig. 4f) as compared to polyQ(7) cells. We next tested whether eIF5A activity is required for ISR-mediated induction of Atf4 using an eIF5A inhibitor, GC7. When added to wild-type cells in the absence of stress, GC7 delayed proliferation (Extended Data Figure 4e). Importantly, under conditions of Tg-induced ER stress, GC7 dampened Atf4 induction (Fig. 4g). Taken together, these experiments indicate that eIF5A activity is required for induction of the translational response to ER stress in striatal cells. We conclude that disruption of eIF5A function in HD dampens the translational response to eIF2a phosphorylation, rendering HD cells more vulnerable to acute stress.

### Reducing translation initiation attenuates ribosome collisions and mitigates ribotoxicity in HD

Given the evidence for ribotoxic stress in HD, we predicted that cells expressing mHTT will be hypersensitized to drugs that induce collisions but benefit from drugs that reduce collisions. Indeed, striatal cells expressing polyQ(111) were hypersensitive to low doses of drugs e.g. emetine (EME) and anisomycin (ANS) that induce ribosome collisions by blocking translation elongation of some but not all elongating ribosomes^50^ (Fig. 4h). By contrast, polyQ(111) cells were less sensitive to low doses of translation initiation inhibitors Rocaglamide A (RocA) and 4EGI-1, targeting initiation factors eIF4A and eIF4E/G1, respectively (Fig. 4h). The distinct and opposing effects of elongation and initiation inhibitors suggest they are not mediated by non-specific reduction of global translation. Instead, we show that such treatments modulate the risk of ribosome collisions. 4EGI-1 increased expression from the collision reporter GFP-K^(AAA)^20, whereas treatment with GC7 to inhibit eIF5A reduced expression (Extended Data Figure 4f). Furthermore, 4EGI-1 decreased and ANS increased premature termination on polyQ(97) in our GFP-HTT-ex1-mCherry reporter (Fig. 4i). Strikingly, 4EGI-1 was previously shown to improve motor symptoms in R6/1 HD mice^51^ and reverse autistic behaviors displayed by eIF4E transgenic mice^52^ supporting a direct link between the molecular events described here and disease etiology. We propose that these beneficial effects in HD models are mediated by reduced risk of collisions, alleviating ribotoxic stress.

## Discussion

Our study provides new mechanistic insights explaining how CAG expansions in mHTT mediate cellular toxicity. We find that the deleterious effects of both mHTT mRNA and protein converge at the level of translation, triggering a cascade of events leading to progressive, system-wide collapse of cellular homeostasis (Fig. 4j). HD-causing mutations give rise to a rapidly decoded tract of glutamines that is immediately flanked by a slowly decoded stretch of prolines, increasing the risk of ribosome collisions. In young healthy cells, robust protein and ribosome quality controls likely resolve and clear collided ribosomes, preventing ribotoxic and proteotoxic stress. As cells age, they become more susceptible to elongation stalls and protein aggregation^41^, exacerbating their ribotoxic effects. As the mutant protein accumulates, it sequesters eIF5A and other RQC factors, leading to profoundly deleterious effects on cellular fitness, including a global decline in proteostasis. Eventually, ribosome pausing and collisions occur throughout the transcriptome, disrupting biogenesis of key proteostasis factors e.g. ribosomes and proteasomes and deregulating polyamine metabolism and stress responses.

Our findings point to a CAG-dependent elongation rate conflict as central to the etiology of HD. Interestingly, CAG repeats—one of the most common homopolymers in eukaryotic proteins—are frequently followed by polyproline stretches encoded by rare codons^53^. This is true for all kingdoms of life, including polyQ tracts in viruses^54^, suggesting a conserved and beneficial use of prolines to buffer the faster elongation on consecutive CAGs^11,55^. Potential benefits of locally reduced elongation rate may include nascent chain folding within the ribosome exit tunnel or engagement of chaperones and other targeting or modifying factors^56,57^. The existence of specialized mechanisms to resolve ribosome slowdown or pausing on polyprolines, as well as the uORF-mediated attenuation of translation initiation, likely mitigate the risk of prolonged ribotoxic collisions on HTT mRNAs with less than 36 CAG repeats. However, expansion of the rapidly decoded tract, combined with a progressive decline in neuronal proteostasis, may overwhelm these homeostatic mechanisms and cause neurotoxicity.

Our model provides a potential mechanistic explanation for several observations in HD. First, the link between CAG repeat length, purity and age of onset^10^: longer expansions will accommodate a larger number of rapidly elongating ribosomes, increasing the risk of collisions and accelerating ribotoxicity. We propose that the presence of less optimal CAA glutamine codons within CAG repeats may reduce collisions by slowing down elongation. Second, ribosome collisions can induce premature termination and release of truncated aggregate-prone mHTT-ex1 fragments in HD^3^. This likely cooperates with missplicing of mHTT intron 1^58^, which yields a fragment that retains the stalling sequence, as well as cleavage of mHTT protein by caspases^59^, to produce neurotoxic mHTT species. The local proximity of mHTT-ex1 nascent chains protruding from neighboring collided ribosomes could even expedite oligomerization and aggregation. Perhaps such a co- or peri-translational mechanism of aggregation may also contribute to the observed depletion of eIF5A. Third, the presence of polyalanine and polyserine protein products in HD brains, thought to occur through ribosome frameshifting on the CAG repeat^60^: given that ribosome collisions induce frameshifting in bacteria^61^ and yeast^62^, it is tempting to speculate they are also responsible for frameshifting on mHTT in mammals.

Of note, ribotoxicity in HD, particularly at later stages, is not limited to translation of mHTT itself. Instead, we find that expression of mHTT is associated with a progressive, age-dependent interference with eIF5A function. In the absence of sufficient eIF5A, elongating ribosomes pause more frequently, leading to collisions that promote nascent chain degradation or aggregation. This generates constant ribotoxic stress that disrupts biogenesis of ribosomes and proteasomes and deregulates polyamine metabolism. Furthermore, we find that loss of eIF5A prevents activation of the pro-survival translational program in response to acute ER stress. Interestingly, mutations affecting eIF5A function are associated with neurodevelopmental disorders in humans^63^, reflecting a potential hypersensitivity of the central nervous system to eIF5A loss. Remarkably, disruption of polyamine metabolism has recently been linked to Parkinson’s Disease^64^, suggesting a possible parallel to other neurodegenerative diseases.

We propose that mHtt-mediated ribosome collisions initiate a cascade of dysfunction that is augmented by eIF5A loss, leading to altered elongation kinetics and a maladaptive response to stress. This progressive cascade impacts central metabolic- and stress-related responses, seeding widespread disruption. Importantly, our findings suggest that pharmacological interventions that reduce ribosome collisions and/or protect eIF5A function could be beneficial and delay onset of symptoms in HD. Remarkably, mild inhibition of translation initiation using 4EGI-1 can reduce ribosome collisions on polyQ-expanded mHTT, providing a mechanistic rationale for the beneficial effects of 4EGI-1 on motor deficits in HD mice^51^. The nuanced and complex interplay between stress, translation initiation and translation elongation in the regulation of mHtt biogenesis uncovered here highlights the importance of improved mechanistic understanding when designing HD therapies.

## Acknowledgments

We thank Dr. Ignacio Munoz-Sanjuan and Dr. Deanna Marchionini, as well as Dr. Kelly Rainbolt and other members of the Frydman lab for their helpful comments. This work was supported by NIH grants GM05643321, NS092525 to JF and AI36178, AI40085, AI091575 to R.An. and Cure Huntington’s Disease Initiative (CHDI) grant A-13887 to JF.

## Author contributions

R.Av. and J.F. designed the study. R.Av. and T.L. carried out experiments, analyzed data and performed statistical analyses. K.H.L performed LC-MS/MS data acquisition. V.B.M and D.G. performed cloning and aggregation experiments. R.Av. and J.F. wrote the manuscript, and all authors approved the manuscript.

## Methods

### Cell cultures

Huh7 human hepatocellular carcinoma cells were grown in DMEM/F-12 1:1 medium (Thermo Fisher) supplemented with 10% fetal bovine serum, 100 units/ml penicillin and 100 mg/ml streptomycin. HEK293 human embryonic kidney cells and STHdh Q7/Q7, Q7/Q111 and Q111/Q111 striatal neuron cells were grown in DMEM high glucose medium (Thermo Fisher) supplemented as above. Cells were grown at 37°C (Huh7, 293T, SH-5YSY) or 32°C (STHdh) in a 5% CO2 incubator.

### Analysis of published ribosome profiling data

gWIPS-viz^65^ and RPFdb^66^ webservers were used to search for relevant ribosome profiling datasets with coverage on Htt uORF and CDS. Relevant mapped reads were downloaded from gWIPS-viz in BigWig format and visualized using Integrated Genome Viewer (IGV) Version 2.8.2.

### Polysome profiles

A total of 1-5×10^7^ striatal neurons or Huh7 cells were harvested by scraping in ice-cold PBS, centrifuged 1000 x g for 5 min at 4°C, resuspended in PBS and centrifuged again. Cell pellets were flash frozen in liquid nitrogen. On the day of experiment, pellets were thawed on ice and resuspended in 200 μl Polysome buffer (25 mM Tris-HCl pH=7.5, 25 mM KCl, 10 mM MgCl_2_, 2 mM dithiothreitol (DTT) and cOmplete EDTA-free protease inhibitor cocktail (Millipore Sigma). Triton X-100 and sodium deoxycholate were added to a final concentration of 1% each and the samples were incubated on ice for 20 min, passed 10 times through a 26G needle and centrifuged at 20,000 x g for 10 min at 4°C to remove cell debris. For RNase-resistance assays 100 U RNase I (Thermo Fisher) was added per 100 μg total RNA in clarified lysates. Digestion was performed for 45 min at room temperature and terminated by addition of 200 U Superase-In (Thermo Fisher). Lysates were loaded on 10-50% sucrose gradients in Polysome buffer and subjected to ultracentrifugation at 36,000 rpm in an SW41.Ti swinging bucket rotor (Beckman Coulter) for 150 min at 4°C. Equal volume fractions were collected using Gradient Station (BioComp) with continuous monitoring of rRNA at UV254.

### Quantitative Real-Time PCR (qRT-PCR)

To extract RNA from sucrose gradient fractions, 1 μl pellet paint co-precipitant (Millipore Sigma) and 500 μl phenol:chloroform:isoamyl alcohol (25:24:1) were added to each 500 μl fraction and incubated for 5 min at room temperature. After centrifugation at 12,000 x g for 15 min, 4°C, the top phase was removed and subjected to another round of extraction as above. 400 μl of the top phase was combined with 600 μl isopropanol and centrifuged at 12,000 x g for 30 min, 4°C. The pellet was washed once with 1 ml 75% ice-cold ethanol, air dried and resuspend in 20 μl RNase-free water. cDNA was synthesized using the High Capacity cDNA Reverse Transcription Kit (Thermo Fisher), according to the manufacturer’s instructions, using 5 μl of RNA from each gradient fraction. qRT-PCR analysis was performed using SensiFast SYBR (BioLine) and gene-specific primers (Supplementary Table 3, plasmids and oligonucleotides), according to the manufacturer’s instructions. To estimate relative abundance of specific mRNAs in each gradient fraction, each Ct value was divided by the sum of Ct values across all gradient fractions.

### SDS-PAGE and immunoblotting

For immunoblotting, adherent cells were washed twice with ice-cold PBS and either lysed on plate with RIPA buffer (25 mM Tris-HCl pH=7.5, 150 mM NaCl, 1% NP-40, 0.5% Sodium deoxycholate) or scraped in PBS, pelleted by centrifugation for 5 min at 1,000 x g, 4°C and resuspended in Polysome buffer. Each lysis buffer was supplemented with fresh 2 mM DTT and Complete EDTA-free protease inhibitor cocktail. RIPA buffer was also supplemented with 50 units/mL benzonase (Millipore Sigma) to remove DNA. For ubiquitination analysis of ribosomal proteins, 100 μM PR-619 and 5 mM NEM (Selleck Chemicals) were included in the lysis buffer. Lysis was performed on ice for 20 min and lysates were clarified by centrifugation for 10 min at 12,000 x g, 4°C. Protein concentration was determined by BCA assay (Thermo Fisher) and 4x Laemmli sample buffer (Bio-Rad) supplemented with fresh 10% 2-mercaptoethanol was added to a final concentration of 1x. 15-30 μg of each sample was resolved on 4-20% (Bio-Rad) or 12% (homemade) SDS-PAGE, transferred to 0.2 or 0.45 μm PVDF membranes presoaked in methanol for 30 sec. Membranes were blocked with 4% molecular biology grade BSA (Millipore Sigma) in tris-buffered saline supplemented with 0.1% Tween-20 (Millipore Sigma) (TBST) for 1 h at room temperature then probed with specific primary antibodies for 2 h at room temperature. Primary antibodies were diluted in 4% BSA/TBST supplemented with 0.02% sodium azide, as follows: rabbit anti-Htt (D7F7, 1:1000); mouse anti-PolyQ (MW1, 1:1000); mouse anti-puromycin (1:2500); mouse anti-eIF2a (1:1000); rabbit anti-p-eIF2a (S51, 1:1000); rabbit anti-RPS2 (1:1000); rabbit anti-RPS3 (1:1000), mouse anti-p38 (1:1000); rabbit anti-p-p38 (T180/Y182, 1:1000); rabbit anti-JNK (1:1000); rabbit anti-p-JNK (T183/T185, 1:1000); rabbit anti-Zak (aa 36-264, 1:500); rabbit anti-Zak (aa 700-750, 1:500), rabbit anti-Atf4 (1:1000), goat anti-Poliovirus VP1 (1:2500); mouse anti-eIF5A (1:1000), rabbit anti-RPL13a (1:2500). Secondary antibodies were diluted 1:10,000 in TBST. Western blot detection was done using ECL Plus Western Blotting Substrate (Thermo Fisher) and images were taken either by film radiography or BioRad GelDoc imager. Alternatively, fluorescent western blot detection was performed using Li-Cor Odyssey Infrared imager. Densitometry analysis was performed using ImageJ version 1.52a.

### Immunofluorescent analysis of Htt aggregation in HEK293 cells

pcDNA3-mClover was generated by PCR amplification and ligation using BamHI and XhoI. Htt-ex1 Q25 or Q97 as described in ^67^ was amplified by PCR and ligated into pcDNA3-mClover using NheI and XhoI. The endogenous 5’UTR of human Htt (NM_001388492.1), either with a functional uORF (wild-type, ATG) or nonfunctional uORF (single nucleotide mutation, AAG), was synthesized by GenScript with flanking BmtI and HindIII sites and ligated into pcDNA3-mClover. For live cell imaging, 3×10^5^ HEK293 cells were seeded in 6 well plates. At 24 h, 2 μg DNA was transfected using Lipofectamine 2000 (Thermo Fisher), according to the manufacturer’s instructions. Transfected cells were monitored once every hour starting from 12 h post-transfection, using a Zeiss Axio Vert.A1 inverted microscope. For aggregate counting, fixed cells were used. 0.8×10^5^ HEK293 cells were seeded on poly-L-ornithine (Millipore Sigma) pre-coated 12 mm coverslips. At 24 h, 400 ng DNA was transfected using Lipofecatmine 2000 (Thermo Fisher), according to the manufacturer’s instructions. Cells were fixed with 4% paraformaldehyde for 15 min at room temperature and washed twice with PBS for 5 min each. Cells were permeabilized with 0.1% triton X-100 for 10 min at room temperature and washed twice with PBS. To label plasma membrane, coverslips were incubated with HCS CellMask Deep Red stain (Thermo Fisher) for 5 min at room temperature and washed twice with PBS. Coverslips were mounted with ProLong Glass Antifade Mountant with NucBlue Stain (Thermo Fisher), dried overnight and stored at −20°C until image acquisition using a Zeiss LSM 700 laser scanning confocal microscope.

### Puromycylation and detection of puromycylated nascent chains

To label and detect nascent chains in cultured cells, 2.5×10^5^ STHdh, Huh7 or SH-SY5Y cells in 6-well plates were incubated with 1 μM puromycin (Thermo Fisher) for 10 min at 32°C (STHdh) or 37°C (Huh7 and SH-SY5Y), washed twice with PBS and lysed on-plate as described above. To label and detect nascent chains in sucrose gradients, each gradient fraction was incubated with 1 μM puromycin for 15 min at 37°C, followed by incubation with 7 μl StrataClean beads (Agilent) for 16 h at 4°C with constant tumbling. Beads were pelleted by centrifugation at 1000 x g for 5 min at 4°C and the supernatant was aspirated. The beads were resuspended in 30 μl 2x Laemmli buffer. Puromycylated samples were resolved on SDS-PAGE as above and monitored using an anti-puromycin antibody (Millipore Sigma).

### Ribosome profiling analysis

Ribosome footprints were prepared essentially as described ^68^. Briefly, adherent cells were washed and harvested in ice-cold PBS by scraping. Cells were pelleted by centrifugation at 1000 x g for 5 min at 4°C and pellets were resuspended in lysis buffer (50 mM HEPES-KOH pH 7.5, 140 mM KCl, 5 mM MgCl2, 0.1% NP-40) supplemented with 0.5 mM DTT and complete EDTA-free Protease Inhibitor Cocktail. Lysates were passed 10 times through a 26G needle and clarified by centrifugation at 20,000 x g for 10 min at 4°C. 25 U RNase I were added per 100 μg total RNA and digestion was performed for 45 min at room temperature, then terminated by addition of 200 U Superase-In (Thermo Fisher). RNase treated samples were fractionated on 10-50% sucrose gradients, as described above, and 80S monosome fractions were collected. RNA was extracted from each fraction using phenol:chloroform:isoamyl alcohol, as described above, and size selected for 16-34 nt using an 8% urea-PAGE. rRNA depletion was performed using Ribo-Zero rRNA Removal Kit (Illumina), according to the manufacturer’s instructions. All downstream steps were performed as described ^68^. Libraries from two independent repeats were sequenced on a HiSeq 4000 (Illumina). After demultiplexing, sequencing reads were trimmed of adaptor sequences and quality filtered using cutadapt (-a CTGTAGGCACCATCAAT -m1 -q20). Ribosomal RNA was removed using Bowtie2 (--un). Remaining reads were aligned to either *Mus musculus* genome assembly GRCm38 (mm10) or poliovirus genome (GenBank V01149.1) using Hisat2 (--trim5 1). Read count matrices were generated using featureCounts and BigWig files were generated using bamCoverage and visualized on Integrated Genome Viewer. For ribosome stall site analysis, the standalone version of PausePred was used ^69^ with the following parameters: window_size=1000, foldchange for pause=20, read_lengths=28,29,30, window coverage%=10, upstream_seq and downstream_seq=25, offset=12,12,12.

### Generation and use of Htt-expressing virus

Htt-ex-1 Q8 or Q73 were amplified by PCR from gWizQ8 or gWizQ73 (kind gift from Donald Lo^70^) using primers with flanking XhoI and EcoRI sites. The primers were designed to amplify the entire ex1 coding region, starting from the CDS AUG and ending with the last codon of ex1, without a stop codon. prib(+)XpA-GFP^71^, encoding for poliovirus type 1 Mahoney with a GFP insert flanked by 2A cleavage sites (2A-GFP-2A), was used as the plasmid backbone. This plasmid has two EcoRI sites: one immediately after the GFP insert, and another in the multiple cloning site (MCS). To delete the latter, the MCS was amplified by PCR using primers with a single nucleotide substitution, and the amplicon was ligated into the backbone using PspOMI and MluI. Htt-ex-1 Q8 or Q73 PCR amplicons were then ligated into the new backbone using XhoI and EcoRI.

To generate viral RNA, 10 μg of the above plasmids were first linearized by incubation with 50 units of MluI for 2 h at 37°C. Linearized DNA was extracted using phenol:chloroform:isoamyl alcohol (25:24:1 v/v), followed by in vitro transcription using MEGAscript T7 Transcription Kit (Thermo Fisher), according to the manufacturer’s instructions. 10 μg of transcribed viral RNA was electroporated into 4×10^6^ Huh7 cells in 0.4 mL PBS, 4 mm electroporation cuvette, at 270 V/960 μF (GenePulser, Bio-Rad). Cells were grown for 24 h at 37°C, at which point culture flasks were subjected to three freeze-thaw cycles and supernatants were centrifuged at 2500 x g for 5 min, 4°C, to remove cell debris. Clarified supernatants containing virus (passage 0, P0) were aliquoted and frozen at −80°C. P0 viruses were amplified once in Huh7 cells to generate P1 virus stock, which was processed and frozen as above. P1 stock was used for all experiments.

To determine virus titers, 1×10^5^ Vero cells were seeded in 6-well plates. The following day, 10-fold serial dilutions of virus stocks/samples were prepared in fresh media without FBS. Cells were washed once with PBS and 0.5 mL inoculum was added for 45 min at 37°C. To prepare the overlay solution, 2% low melting point agarose (Thermo Fisher) was dissolved in water by microwaving and combined 1:1 v/v with 2x media supplemented with 2% FBS just before use. 2 ml overlay solution was added to each well and allowed to gel at room temperature. Two days after infection, 2 ml 2% formaldehyde was added for 1 h at room temperature. Formaldehyde and overlay were removed and cells were stained with 1 ml 0.1% crystal violet in 20% ethanol for 1 h at room temperature. Plates were disinfected using 1% bleach and plaques were counted. For infections, Huh7 or SH-SY5Y cells were washed once with PBS and virus stock, at a multiplicity of infection (MOI)=10, was added to fresh media without FBS. After 45 min at 37°C, inoculum was replaced with fresh media supplemented with 10% FBS.

### Analysis of published proteomic datasets

Processed mass-spec intensity tables were downloaded from the supplementary information section of the following published articles: ^6^, Table S1 and S3 (label-free quantification, LFQ);^38^, Table S1 and S3 (LFQ); ^36^, Table S2 (LFQ); ^37^, Table S1 (raw intensities). No additional normalization of the data was performed. For^6^ dataset, separate areas of the brain were plotted as independent datapoints. Statistical tests were done using two-sided Student’s t-test.

### Generation and extraction of urea-resistant aggregates from transfected HEK293 cells

Htt-ex1 Q25 or Q97 as described in ^67^ was amplified by PCR to include a C-terminal HA tag and ligated into pcDNA3 using NheI and XhoI. 1.8 x 10^7^ HEK293 cells were plated on gelatin-coated 15 cm plates. The following day, cells were transfected with 30 μg pcDNA3-Htt-ex1-Q25-HA or pcDNA3-Htt-ex1-Q97-HA using Lipofectamine 2000 (Thermo Fisher), according to the manufacturer’s instructions. At 30 h post-transfection, cells were washed twice with ice cold PBS and scraped into the same buffer. Cells were pelleted by centrifugation at 800 x g for 5 min at 4°C and washed again with the same buffer. Pellets were resuspended in 1 ml lysis buffer (100 mM NaCl, 50 mM HEPES pH 7.4, 0.5% NP-40, 1 mM PMSF, Benzonase and Complete protease inhibitors) and lysed using tip sonication. Lysates were clarified by gentle centrifugation at 800 x g for 2 min at 4°C. Supernatant was collected and protein concentration was determined using the BCA kit (Thermo Fisher). Clarified lysates were adjusted to a final volume of 800 μl at equivalent concentrations (3.4 mg/ml) using lysis buffer. Insoluble fractions were pelleted by centrifugation at 21,000 x g for 30 min at 4°C. Supernatant was removed and pellets were resuspended in a strong wash buffer (8 M urea, 100 mM NaCl, 100 mM HEPES pH 7.4) to remove soluble proteins from aggregates. Insoluble material was pelleted and washed again as above. Remaining pellets were resuspended in 200 μl protein buffer (50 mM HEPES, 100 mM NaCl) by sonication. Formic acid was then added to a final concentration of 90% to dissolve remaining aggregates and dried using a SpeedVac for downstream proteomic analysis.

### Sample preparation for proteomic analysis

Proteins were extracted from isolated polysomes or aggregates using methanol-chloroform precipitation. 400 μl methanol, 100 μl chloroform and 350 μl water were added sequentially to each 100 μl sample, followed by centrifugation at 14,000 x g for 5 min at room temperature. The top phase was removed and the protein interphase was precipitated by addition of 400 μl methanol, followed by centrifugation at 14,000 g for 5 min at room temperature. Pellet was air dried and resuspended in 8M urea, 25 mM ammonium bicarbonate (pH 7.5). Protein concentration was determined by BCA (Thermo Fisher) and 2-4 μg total protein were subjected to reduction and alkylation by incubation with 10 mM DTT for 1 h at room temperature followed by 5 mM iodoacetamide for 45 min at room temperature, in the dark. The samples were then incubated with 1:50 enzyme to protein ratio of sequencing-grade trypsin (Promega) overnight at 37 °C. Peptides were desalted with μC18 Ziptips (Millipore Sigma), dried and resuspended in 10 μL 0.1% formic acid in water.

### LC-MS/MS acquisition

LC-MS/MS analyses were conducted using either a QExactive Plus Orbitrap (QE, RNase-digested polysomes) or a Velos Pro Elite Orbitrap (Elite, virus polysomes and HEK293 aggregates) mass spectrometer (Thermo Fisher) coupled online to a nanoAcquity UPLC system (Waters Corporation) through an EASY-Spray nanoESI ion source (Thermo Fisher). Peptides were loaded onto an EASY-Spray column (75 μm x 15 cm column packed with 3μm, 100 Å PepMap C18 resin) at 2% B (0.1% formic acid in acetonitrile) for 20 min at a flow rate of 600nl/min. Peptides were separated at 400nL/min using a gradient from 2% to 25% B over 48 min (QE) or 220 min (Elite) followed by a second gradient from 25% to 37% B over 8 minutes and then a column wash at 75% B and reequilibration at 2% B. Precursor scans were acquired in the Orbitrap analyzer (QE: 350-1500 m/z, resolution: 70,000@200 m/z, AGC target: 3e6; Elite: 300-1800 m/z, resolution: 60,000@400 m/z, AGC target: 2e6. The top 10 or 6 (QE or Elite) most intense, doubly charged or higher ions were isolated (4 m/z isolation window), subjected to high-energy collisional <u>dissociation</u> (QE: 25 NCE; Elite: 27.5 NCE), and the product ions measured in the Orbitrap analyzer (QEplus resolution: 17,500@200 m/z, AGC target: 5e4; Elite resolution: 15,000@400 m/z, AGC target: 9e4).

### Mass spectrometry data processing

Raw MS data were processed using MaxQuant version 1.6.7.0 ^72^. MS/MS spectra searches were performed using the Andromeda search engine ^73^ against the forward and reverse human and mouse Uniprot databases (downloaded August 28, 2017 and November 25, 2020, respectively). Cysteine carbamidomethylation was chosen as fixed modification and methionine oxidation and N-terminal acetylation as variable modifications. Parent peptides and fragment ions were searched with maximal mass deviation of 6 and 20 ppm, respectively. Mass recalibration was performed with a window of 20 ppm. Maximum allowed false discovery rate (FDR) was <0.01 at both the peptide and protein levels, based on a standard target-decoy database approach. The “calculate peak properties” and “match between runs” options were enabled.

All statistical tests were performed with Perseus version 1.6.7.0 using either ProteinGroups or Peptides output tables from MaxQuant. Potential contaminants, proteins identified in the reverse dataset and proteins only identified by site were filtered out. Intensity-based absolute quantification (iBAQ) was used to estimate absolute protein abundance. Two-sided Student’s t-test with a permutation-based FDR of 0.01 and S0 of 0.1 with 250 randomizations was used to determine statistically significant differences between grouped replicates. Categorical annotation was based on Gene Ontology Biological Process (GOBP), Molecular Function (GOMF) and Cellular Component (GOCC), as well as protein complex assembly by CORUM.

### Cell viability

1×10^4^ STHdh cells were seeded per well in a black wall, clear bottom 96-well plate. At 24 h, thapsigargin (Tg, AG Scientific) was added for another 24 hours. Cell viability was assayed using CellTiter-Glo (Promega), according to the manufacturer’s instructions. Luminescense was recorded using a BMG Labtech CLARIOstar^®^ Microplate Reader.

### Polyamine Quantification

Polyamine concentration was measured using Total Polyamine Assay Kit (Abcam), according to the manufacturer’s instructions. 2×10^6^ STHdh striatal neurons were harvested by scraping in ice-cold PBS and resuspended in 100 μl ice-cold polyamine assay buffer. Samples were homogenized on ice using a Dounce homogenizer and clarified by centrifugation at 10,000 x g for 5 min at 4°C. 2 μl sample clean-up buffer was added to each sample and incubated for 30 min at room temperature. Sampled were passed through a 10 kD MWCO filter (Thermo Fisher) by centrifugation at 10,000 x g for 10 min. 5 μl filtrate (each) was used for background test, polyamine test and BCA protein quantification (Thermo Fisher). Reaction buffer was added for 30 mins at 37°C, protected from light. Fluorescence (Ex/Em=535/587 nm) was detected using a BMG Labtech CLARIOstar® Microplate Reader. Total polyamine concentration was calculated by subtracting the background value and normalizing based on total protein.

### Data availability

Sequencing data were deposited in SRA database under BioProject number PRJNA730032. The mass spectrometry proteomics data have been deposited to the ProteomeXchange Consortium via the PRIDE^74^ partner repository with the dataset identifier PXD026012.

---

**Extended data figure 1. uORF decreases HTT translation under normal condition and increases it during stress. (a)** Position and amino acid sequence of mammalian HTT uORF and start of CDS. **(b)** Ribosome profiling traces of initiating ribosomes (A-sites, top) and elongating ribosomes (footprints, bottom) on human HTT uORF and start of CDS. (c) 293T cells were transfected with constructs expressing wild-type polyQ(25) or mutant polyQ(97) HTT-ex1 with a C-terminal fluorescent GFP. Shown are live cell images at the indicated times. (d) Top, ribosome profiling traces of elongating ribosomes on Htt uORF and start of CDS in different organisms and tissues. Bottom, Htt protein level (low, moderate, high) in the indicated tissues, from ProteinAtlas. **(e)** Wild-type mouse striatal cells were treated with 1uM Tg for 1h and subjected to polysome profile analysis on sucrose gradients. RNA was extracted from gradient fractions and analyzed by qPCR to measure the indicated transcripts. RNA levels in each fraction were calculated as percent of total across all gradient fractions. Shown are means +/-s.d. of 3 independent repeats.

**Extended data figure 2. Translation of mHTT from poliovirus genome is associated with ribosome collisions and premature termination. (a)** Poliovirus replicates more slowly when engineered to express mutant versus wild-type HTT-ex1. Wild-type polyQ(8) and mutant polyQ(73) human HTT-ex1 were cloned into poliovirus genome. Infectious viruses were generated by transfection of *in vitro* transcribed RNA. Plaque assays were performed cells to determine virus titers. Shown are means +/-s.d.of 3 independent repeats (left) and representative images of viral plaques (right). **(b-c)** Huh7 cells (b) and SH-SY5Y neuroblasts (c) were infected with engineered viruses. At the indicated times, puromycin was added to tissue culture media to label nascent polypeptide chains and lysates were analyzed by immunoblotting. Shown is a representative of 2 independent repeats. Purple boxes indicate virus-Htt polyproteins prior to cleavage by the viral protease. **(d)** Huh7 cells were infected as before, lysed and treated with RNase I. Lysates were fractionated on 10-50% sucrose gradients and fractions corresponding to 80S monosomes, disomes and trisomes were incubated with puromycin, followed by immunoblot analysis. Shown is a representative of 3 independent repeats. **(e)** Huh7 cells were infected as before and subjected to polysome profile analysis on 10-50% sucrose gradients. Puromycin was added to gradient fractions to label nascent chains, followed by immunoblot analysis. Shown are representative blots of 2 independent repeats. **(f)** Huh7 cells were infected as before, lysed and treated with RNase I. Lysates were fractionated on 10-50% sucrose gradients and 80S fractions were collected for ribosome profiling analysis (n=2). Shown are relative ribosome densities on viral sequences either upstream of or downstream to the HTT-ex1 insert.

**Extended data figure 3. Evidence of ribotoxicity in HD models. (a-b)** Levels of RQC factors increase, while those of elongation factor decrease, in the soluble brain proteome of presymptomatic R6/2 HD mice. Shown are absolute levels of the indicated proteins in wild-type and R6/2 HD cells at 5, 8 and 12 weeks of age (a) and their relative differences between wild-type and R6/2 at 8 weeks (b). **(c)** Pairwise comparison of individual protein levels, as detected by MS, in the insoluble brain proteome of wild-type and zQ175 HD mice at 40 weeks of age. **(d)** Relative ubiquitination of specific sites on ribosomal proteins, not affected by collisions, as determined by MS analysis of soluble brain proteome in wild-type and zQ175 mice at 40 weeks of age. **(e)** Ribosome profiling traces of mRNAs showing increased pausing in striatal cells expressing polyQ(111). **(f)** Stoichiometry of ribosomal (top) and proteasomal (bottom) proteins is disrupted in R6/2 brains. Shown are kurtosis scores for all core ribosomal or proteasomal proteins as measured by MS.

**Extended data figure 4. Aberrant stress translation in HD striatal cells and reversal by small molecule inhibitors. (a)** Global translation is lower in striatal cells expressing two copies of mHtt polyQ(111). Translation rates were monitored by puromycin labeling and quantified by densitometry (n=3). **(b)** Translation of ISR-responsive transcripts is lower in striatal cells expressing mHtt polyQ(111). Shown are pairwise comparison of ribosome densities (as RPKM) from ribo-seq experiments (n=2). **(c)** Htt uORF bypassing is reduced in striatal cells expressing mHtt polyQ(111). **(e)** Striatal cells were treated with different concentrations of GC7 to inhibit eIF5A. At 48h, the difference in proliferation from 24h was measured by CellTiterGlo (n=4). **(f)** Wild-type striatal cells were transfected with plasmids encoding GFP upstream to a K^(AAA)^20 ribosome collision/termination motif. Drugs were added 6h post-transfection, as indicated. Percent GFP-positive cells was measured using flow cytometry at 24h post-transfection (n=3).

