## Supplementary figures and images for "Ribotoxic collisions on CAG expansions disrupt proteostasis and stress responses in Huntington’s Disease"

### Extended Data Figures

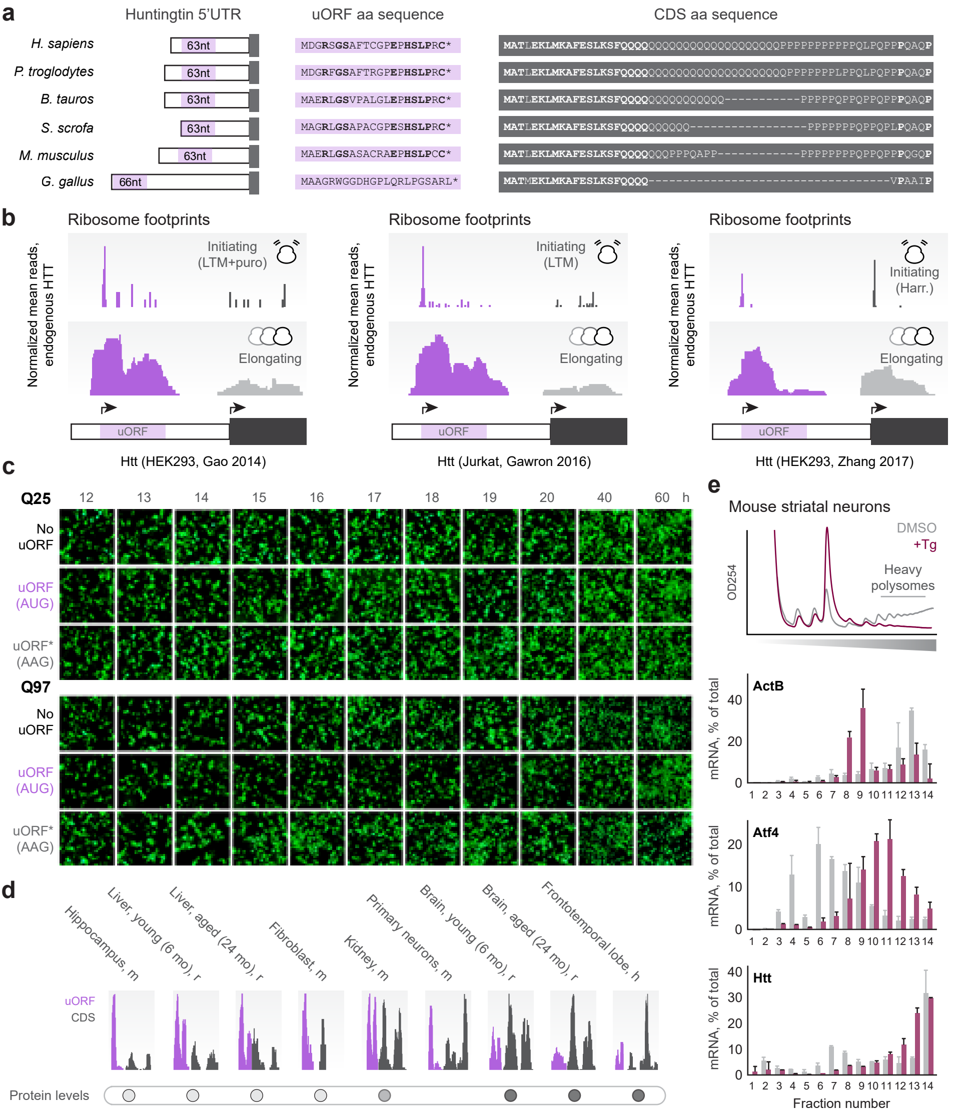

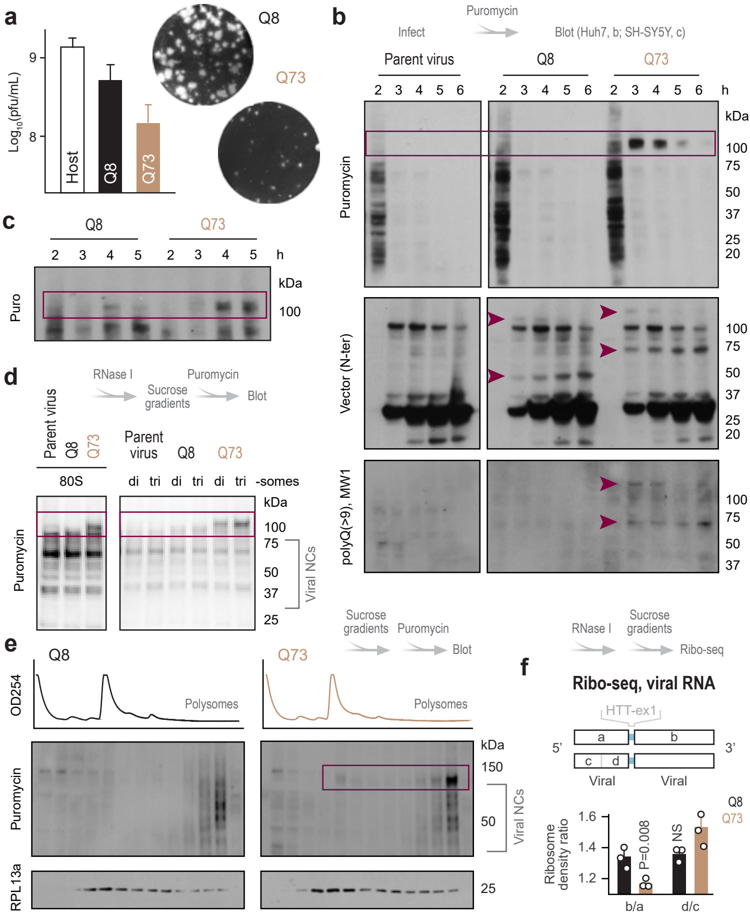

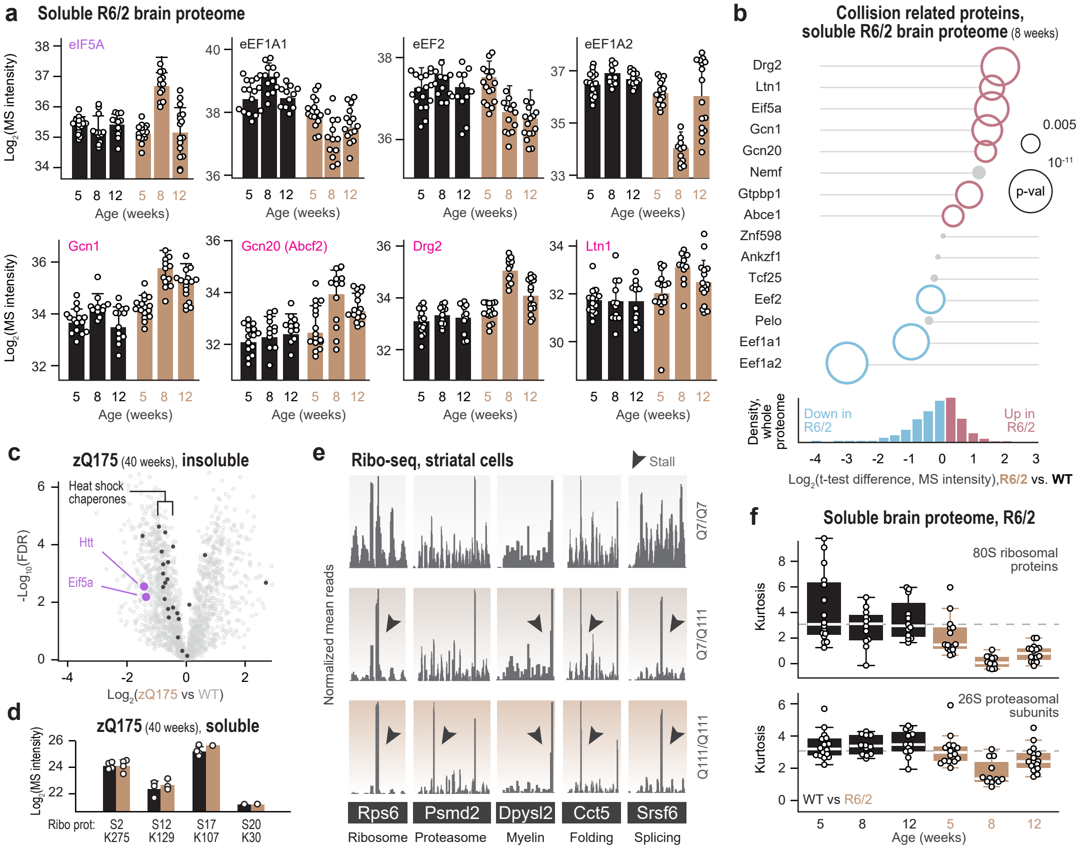

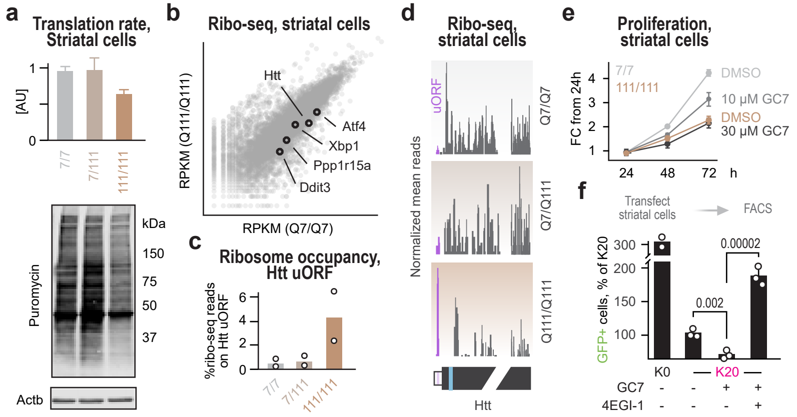
